# MAF1 is a Chronic Repressor of RNA Polymerase III Transcription in the Mouse

**DOI:** 10.1101/775353

**Authors:** Nicolas Bonhoure, Viviane Praz, Robyn D. Moir, Gilles Willemin, François Mange, Catherine Moret, Ian M. Willis, Nouria Hernandez

**Affiliations:** Center for Integrative Genomics, Faculty of Biology and Medicine, University of Lausanne, 1015 Lausanne, Switzerland; Swiss Institute of Bioinformatics, 1015 Lausanne, Switzerland; Department of Biochemistry, Albert Einstein College of Medicine, Bronx, New York 10461, USA; Department of Systems and Computational Biology, Albert Einstein College of Medicine, Bronx, New York 10461, USA

## Abstract

*Maf1*^*-/-*^ mice are lean, obesity-resistant and metabolically inefficient. Their increased energy expenditure is thought to be driven by a futile RNA cycle that reprograms metabolism to meet an increased demand for nucleotides stemming from the deregulation of RNA polymerase (pol) III transcription. Metabolic changes consistent with this model have been reported in both fasted and refed mice, however the impact of the fasting-refeeding-cycle on pol III function has not been examined. Here we show that changes in pol III occupancy in the liver of fasted versus refed wild-type mice are largely confined to low and intermediate occupancy genes; high occupancy genes are unchanged. However, in *Maf1*^*-/-*^ mice, pol III occupancy of the vast majority of active loci in liver and the levels of specific precursor tRNAs in this tissue and other organs are higher than wild-type in both fasted and refed conditions. Thus, MAF1 functions as a chronic repressor of active pol III loci and can modulate transcription under different conditions. Our findings support the futile RNA cycle hypothesis, elaborate the mechanism of pol III repression by MAF1 and demonstrate a modest effect of MAF1 on global translation via reduced mRNA levels and translation efficiencies for several ribosomal proteins.

## Introduction

The synthesis of the translational apparatus is energetically costly and, consequently, subject to tight regulation (see ^1^ and references therein). For example, the activities of both pol I and III are controlled in response to nutrient availability ^2,3–5,6^. In the case of pol III, this control is exerted largely by the MAF1 protein, which constitutes its main repressor ^7^. In yeast, under favorable growth conditions, phosphorylation of Maf1 in a TOR complex 1- (TORC1) and protein kinase A-dependent manner inhibits its interaction with pol III and results in its nuclear exclusion ^4,5^. Upon nutrient deprivation or stresses like DNA damage, TORC1 is inactivated, Maf1 becomes dephosphorylated and shuttles into the nucleus where it represses pol III transcription. In mammalian cells, mTORC1 directly phosphorylates MAF1 on three residues (S60, S68, and S75)^8^. Following serum deprivation, MAF1 undergoes dephosphorylation, increasingly targets pol III genes as determined by DamIP-seq, and represses pol III transcription ^9,10^.

Mice lacking *Maf1* are viable and fertile. They are slightly smaller and leaner than their wild-type (WT) counterparts, and they are resistant to diet-induced obesity and non-alcoholic fatty liver disease ^11^. This is due in part to reduced food intake but also to reduced metabolic efficiency. *Maf1*^*-/-*^ mice display increased energy expenditure throughout the diurnal cycle even though they are not physically more active than WT mice ^11^. Studies with fasted mice revealed elevated pol III transcription and precursor tRNA levels in the liver and numerous others tissues (∼3-9 fold changes in precursor tRNAs) but no change in total tRNA or mature tRNA levels. These and other findings led to the proposal of a futile RNA cycle to account for the lean phenotype and wasteful use of metabolic energy in *Maf1*^*-/-*^ mice ^11^. A key feature of this hypothesis is the existence of a homeostatic mechanism for pol III transcripts, most notably mature tRNAs, that keeps their cellular levels largely constant when pol III transcription is increased by deletion of *Maf1*. Important predictions of the futile RNA cycle hypothesis have recently been confirmed by targeted metabolomics ^12^. The findings include changes in liver metabolites involved in de novo nucleotide synthesis and RNA turnover and changes in pyruvate and acetyl CoA at the interface of glycolysis and the TCA cycle. In the context of a mouse model of increased energy demand, these findings are further supported by increased lipolysis in white adipose tissue (WAT) and increased autophagy/lipophagy in the liver, processes that together provide TCA cycle substrates for enhanced energy metabolism ^13^. Metabolomic support for changes in these processes has also be obtained: The levels of many amino acids are increased in liver and skeletal muscle and changes in acyl carnitine levels are consistent with increased oxidation of fatty acids ^11,12^. Importantly, while a futile RNA cycle may be the main driver of energy expenditure in *Maf1*^*-/-*^ mice, other energy costs coming from enhanced cycling of hepatic lipids and increased activity of the urea cycle are likely incurred indirectly and contribute to the overall energy budget ^11,12^. Consistent with energy expenditure being higher in *Maf1*^*-/-*^ mice at night (i.e. when mice are actively feeding), changes in amino acid and fatty acid metabolism are more pronounced in refed versus fasted animals ^12^. Thus, somewhat counterintuitively (since MAF1 function is inhibited by nutrient signaling), futile RNA cycling is likely to be higher in the refed state. This issue has yet to be examined.

The smaller size and the lean phenotype of *Maf1*^-/-^ mice is notably different from the phenotypes observed following perturbations of MAF1 expression in *D. melanogaster* ^14^ and *C. elegans* ^15^. In the fly, a decrease in MAF1 levels leads to increased larvae volume, and deletion of *Maf1* specifically in the fat body of fly larvae increases mature tRNA levels, which in turn leads to increased translation. In *C. elegans*, increasing or decreasing MAF1 levels causes reciprocal changes in body area. Strikingly, in worms and in cultured mammalian cells, perturbation of MAF1 levels correlates with changes in >1000 mRNAs ^15,16^. Common effects of MAF1 on the expression of fatty acid synthase (*Fasn*) and acetyl CoA carboxylase alpha (Acaca) have been found in worms and mammalian cells and is correlated with changes in fat accumulation ^15,17^. Additionally, recent work indicates that MAF1 is important for the normal function of mouse embryonic stem cells and in cell fate determination. Specifically, loss of MAF1 was shown to impair adipogenesis ^16^. In previous studies we profiled the pol II transcriptome in white adipose tissue of WT and *Maf1*^-/-^ mice and found no significant differences ^11^. However, the studies noted above suggest that a wider examination of the effect of MAF1 on mouse gene expression is warranted.

In this study, we used wild-type and *Maf1*^-/-^ mice to examine the role of MAF1 in pol III transcription during the fasting-feeding cycle. We found that pol III occupancy in the liver of WT mice increases during the fasted to refed transition but this response is limited to mostly low and intermediate occupancy genes; high occupancy genes are essentially unchanged. Similar findings involving overlapping subsets of stable and changing genes are seen following a partial hepatectomy ^18^. In contrast, in *Maf1*^*-/-*^ mice, pol III occupancy was increased for the majority of active loci in liver, regardless of their occupancy level, and the levels of specific precursor tRNAs in this tissue and in other organs are higher than wild-type in both fasted and refed conditions. Thus, MAF1 keeps pol III transcription in check in different metabolic states. We also report changes in the pol II transcriptome in refed liver and a modest reduction in global translation in the refed state that reflects decreased mRNA levels and/or translation efficiencies of certain ribosomal proteins.

## Results

### Fasting and refeeding affects pol III occupancy in mouse liver

In response to feeding and fasting, mammalian metabolism switches between the utilization of carbohydrates and that of fats as principal metabolic fuels to maintain glucose homeostasis and cellular function. In the process, changes in hormonal and nutrient signalling lead to extensive reprogramming of gene expression in many tissues, especially the liver ^19^. Changes in nutrient signalling have been shown to impact pol III gene regulation in lower eukaryotes, invertebrates and in cultured mammalian cells ^10,20^. However, a detailed study of the effect of fasting and refeeding on pol III transcription in mice has not been reported. To assess this, we first measured genome-wide pol III occupancy by chromatin immunoprecipitation and high throughput sequencing (ChIP-seq) as a proxy for pol III transcription ^9,21^. We compared liver samples from wild-type mice fasted for 8 hours with samples from mice that had been fasted and refed for 4 hours. As expected, the median pol III occupancy score decreased after fasting (Figure 1A). However, this result primarily reflected lower scores for genes in the bottom three quintiles (yellow, light-green, and dark-green dots) (Figure 1A). Genes scoring in the top two quintiles generally remained highly occupied after fasting. Limma analysis found more than one hundred loci with significantly lower occupancy scores after fasting (Figure 1B, red dots); these included nearly all 5S genes as well as many tRNA genes and SINEs. In contrast, the 4.5S genes, pol III genes with type 3 promoters including the expressed selenocysteine tRNA gene, and a subset of tRNA genes encompassing all amino acid isotypes were similarly occupied in both conditions (Figure 1B, black dots, see Tables S1 and S2). The preceding observations and parallel findings in serum starved human fibroblasts ^9^ indicate that active pol III loci can be partitioned into two groups based on pol III occupancy; one that is sensitive to nutrient signals and one that is not very sensitive to these signals.

**Figure 1.**
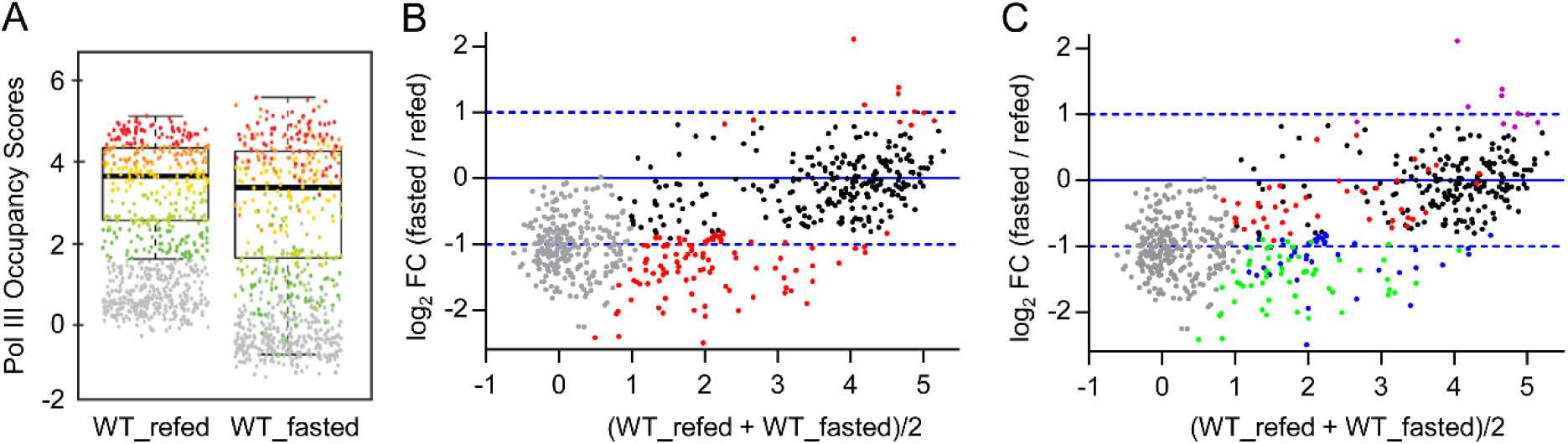
Altered pol III occupancy in livers from fasted and refed WT mice. (A) Average pol III occupancy scores, Log2(IP/Input), for two replicate ChIP-seq experiments performed with pooled liver samples from three wild-type mice fasted for 8 h or fasted for 8 h and refed for 4 h. Each dot corresponds to a pol III locus and is colored according to its score quantile in the WT_Refed sample. (B) MvA plot showing the average occupancy scores in refed and fasted conditions (x-axis) and the log2 fold change (fasted/refed) between these conditions (y-axis). Red dots, loci with significant changes by limma analysis (adj. p-value < 0.01); black dots, pol III loci with scores above the cut-off in at least one sample that were not significant by limma analysis (adj. p-value ≥ 0.01), grey dots, pol III loci with scores below the cut-off in both samples. See Table S2 for corresponding scores and limma analysis. (C) Color mapping of pol III occupancy relationships between nutritional and proliferative signaling responses. The fasting/refeeding data in panel B are colored based on the fold change in occupancy scores before and 36 hours after partial hepatectomy (PH)^18^. Black dots, no occupancy change upon feeding/fasting or PH; green dots, reciprocal changes in pol III occupancy upon feeding/fasting and PH; red dots, genes with higher sensitivity to PH than to feeding; blue and magenta dots, genes with higher sensitivity to feeding than to PH.

Genome-wide changes in pol III occupancy have recently been reported in mouse liver following partial hepatectomy (PH) ^18^. This treatment results in the synchronous entry of hepatocytes into the cell cycle during liver regeneration. We therefore explored possible relationships between genes whose pol III occupancy changed, or not, in response to nutritional versus proliferative signals. We found significant overlap between the changing and stably occupied genes under both conditions: (i) Of the 212 high occupancy genes in the PH dataset (average occupancy score > 3.5, see Table S4 in ref ^18^), 197 were not significantly changed by PH. Of these, 90% (177 genes) were also not significantly changed upon fasting (Figure 1C, black dots). Thus, pol III occupancy of these genes is stable in the face of changing nutritional and proliferative signals. (ii) Of the 143 genes with significantly increased occupancy after PH, 55 genes showed a reciprocal response upon fasting (Figure 1C, green dots), demonstrating that these genes are sensitive to both nutritional and proliferative signals. The responses of other genes in the feeding/fasting and PH datasets suggest differential sensitivities to the respective signals: among the genes with significantly higher occupancy after PH were 40 genes with low to medium occupancy scores that were unchanged upon fasting (Figure 1C, red dots). These genes appear to be more responsive to proliferative than nutritional signals. Conversely, 61 genes with significantly lower pol III occupancy after fasting were unchanged after PH (Figure 1C, blue dots). This group contained 30 out of 34 active 5S rRNA genes suggesting more robust pol III control of ribosome synthesis by nutrients than by PH. Overall these data indicate that pol III occupancy in wild-type mouse liver can be stable or vary depending on the gene and on cell physiology.

### Higher pol III occupancy in *Maf1*^-/-^ mouse liver

We next determined the contribution of MAF1 to pol III occupancy in the fasting and refeeding regime. Out of 480 loci bound by pol III in WT or *Maf1*^-/-^ mouse liver in either condition, 425 loci showed significant occupancy differences between WT and *Maf1*^-/-^ samples: 257 showed higher pol III occupancy in the knockout in both the refed and the fasted state, only 25 loci were more highly occupied only under the fasted condition, and 143 were more highly occupied only under the refed condition (Figure 2A). In addition, the score differences between WT and *Maf1*^-/-^ samples were larger in the refed condition than in the fasted condition (Figure 2B and 2C, Tables S3 and S4). These data suggest that MAF1 represses pol III transcription in both fasted and refed conditions, and show that remarkably, the quantitative effect of its deletion is more pronounced in the refed state. Accordingly, in *Maf1*^-/-^ mice, pol III occupancy scores were significantly decreased in fasted relative to refed conditions (Figure 2D and Table S5). These findings are consistent with our previous observation that MAF1 represses pol III occupancy of its target genes in cultured human fibroblasts under both repressive serum-starved conditions and favorable serum-replete conditions ^9^. Moreover, the data suggest the existence of MAF1-independent mechanisms that regulate pol III occupancy in mouse liver during the fasting-feeding cycle.

**Figure 2.**
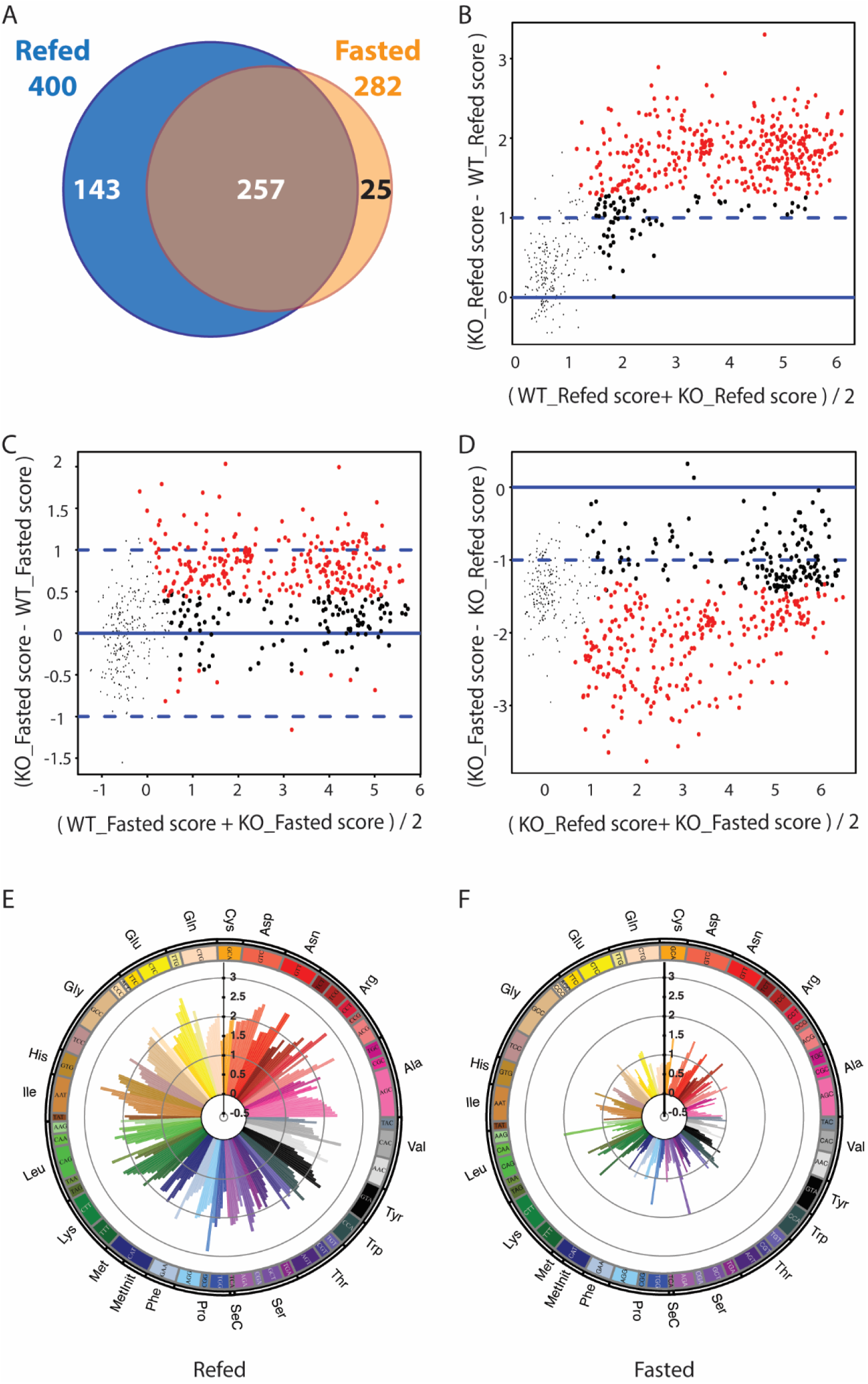
Differential effects of *Maf1* loss on pol III occupancy in mouse liver upon fasting and refeeding. (A) Venn diagram showing the number of pol III loci with significantly higher pol III occupancy in *Maf1*^-/-^ versus WT liver samples in the fed or fasted condition (adj. p-value ≤ 0.01). (B) MvA plot comparing liver samples from refed WT and *Maf1*^-/-^ mice. Red dots, loci with significant changes by limma analysis (adj. p-value < 0.01); large black dots, pol III loci with scores above the cut-off in at least one sample that were not significant by limma analysis (adj. p-value ≥ 0.01), small black dots, pol III loci with scores below the cut-off in both samples. See Table S3 for scores and limma analysis. (C) MvA plot comparing liver samples from fasted WT and *Maf1*^-/-^ mice. Dots are colored as in panel B. See Table S4 for scores and limma analysis. (D) MvA plot comparing liver samples from refed and fasted *Maf1*^-/-^ mice. Dots are colored as in panel B. See Table S5 for scores and limma analysis. (E) Circular plot indicating the positive pol III occupancy score ratios (*Maf1*^-/-^ versus WT) for each annotated tRNA gene occupied in at least one sample in the refed condition. (The few genes with negative scores are not shown to avoid compression of the fold change scale). (F) As in panel E but for the fasted condition.

The 257 loci whose occupancy was higher in the *Maf1*^-/-^ samples under both fasted and refed conditions included most Rn5s (5S) genes, three Rn4.5s (4.5S) genes and several genes with diverse functions (“other pol III genes” in Tables S1-S5, specifically Rny1, RnU6atac, RnU6, Rmrp, and Rn7sl genes), about 30 SINEs and 172 tRNA genes, i.e. 60% of the 288 tRNA genes occupied in at least one genotype and one condition. Most Rn4.5s genes and 92 tRNA genes showed increased pol III occupancy in the *Maf1*^-/-^ samples specifically in the refed condition. Given the potential for differential tRNA gene expression to affect translation, we examined positive score differences in *Maf1*^-/-^ versus WT liver for all occupied tRNA genes and organized these data by isoacceptors (Figures 2E and 2F) and isotypes (Figures S1C and S1D). The score differences were generally larger in the refed state (Figures 2E and 2F), consistent with the results above, but varied from gene to gene within each isoacceptor class. We then compared the cumulative scores for all tRNA genes for each of the 22 isotype classes (tRNA^Met^ genes were considered separately from tRNAi^Met^ genes) and found that the increase in pol III occupancy associated with the absence of MAF1 was general; the various isotypes remained in the same quartile in the *Maf1*^-/-^ samples, both under refed and fasted conditions (Figures S1A and S1B). However, isotypes in the more occupied quantiles displayed somewhat larger differences in pol III occupancy than less occupied quantiles, as shown in Figures S1C and S1D, where isotypes were ordered according to their occupancy scores in the WT samples. This effect was especially marked in the refed samples (Figure S1C), and was also noticeable with scores cumulated by isoacceptors rather than by isotypes (Figure S1E and F). Thus, for tRNA genes occupied by pol III in WT or *Maf1*^-/-^ liver in either condition, the absence of MAF1 results generally in higher pol III occupancy. In contrast, many of the annotated tRNA genes that were unoccupied by pol III in WT liver (198 out of 433 annotated tRNA genes) remained so in the *Maf1*^-/-^ liver samples (148 out of 433), confirming that as in human IMR90Tert cells ^9^, transcription from a large percentage of tRNA genes is stably repressed.

### *Maf1*^-/-^ mice have higher levels of pre-tRNAs in fasted and refed conditions

Previous studies with fasted *Maf1*^-/-^ liver found increased pol III occupancy and precursor tRNA levels but no significant change in mature tRNA levels ^11,22^. To determine whether or how these findings might change given the greater effect of MAF1 on pol III occupancy after refeeding, we examined the levels of several intron- and non-intron-containing precursor tRNAs and mature tRNAs in liver and various other tissues from fasted and refed mice by northern analysis. Comparing *Maf1*^-/-^ and WT liver samples, intron-containing pre-tRNA^Ile^-TAT, pre-tRNA^Leu^-CAA and pre-tRNA^Tyr^-GTA increased from ∼30-80% in the fed *Maf1*^-/-^ samples (Fig. 3A, left panel and Figure 3C). In contrast, the levels of the corresponding mature tRNAs were unchanged (Figure 3A, left panel and Figure 3D). Similar increases of 32% and 54% were observed for the non-intron-containing precursors, pre-tRNA^Ser^-CGA and pre-tRNA^His^-GTG, in the fed *Maf1*^-/-^ samples and the corresponding mature tRNAs along with mature tRNA^Glu^-CTC and tRNA^Leu^-AAG were again unchanged (Figure 3A, left panel and Figure 3D). Curiously, the overall magnitude of effect of *Maf1*^*-/-*^ on the five precursor tRNAs was considerably lower in refed liver than in fasted liver where northern analysis measured three-fold changes for the same intron-containing pre-tRNAs (Figure 3B) ^11^, an observation that does not correlate with the higher pol III occupancy on these genes in the refed state in this tissue (Table S1). In contrast to the less than two-fold changes for precursor tRNAs in refed liver, all other tissues from *Maf1*^*-/-*^ mice showed robust increases in precursor tRNA levels in both fasted and refed conditions (Figure 3A, right panel, Figure 3B) while in WT tissues, precursor tRNA levels were uniformly reduced upon fasting. Thus, pol III occupancy and RNA analyses indicate that fasting and refeeding affect pol III transcription in WT animals, although the magnitude of the effect varies in different tissues, and that *Maf1*^-/-^ animals have higher pol III occupancy and transcription in both the fasted and refed conditions as compared to WT animals. Accordingly, MAF1 functions to limit pol III transcription in both fasted and refed conditions.

**Figure 3.**
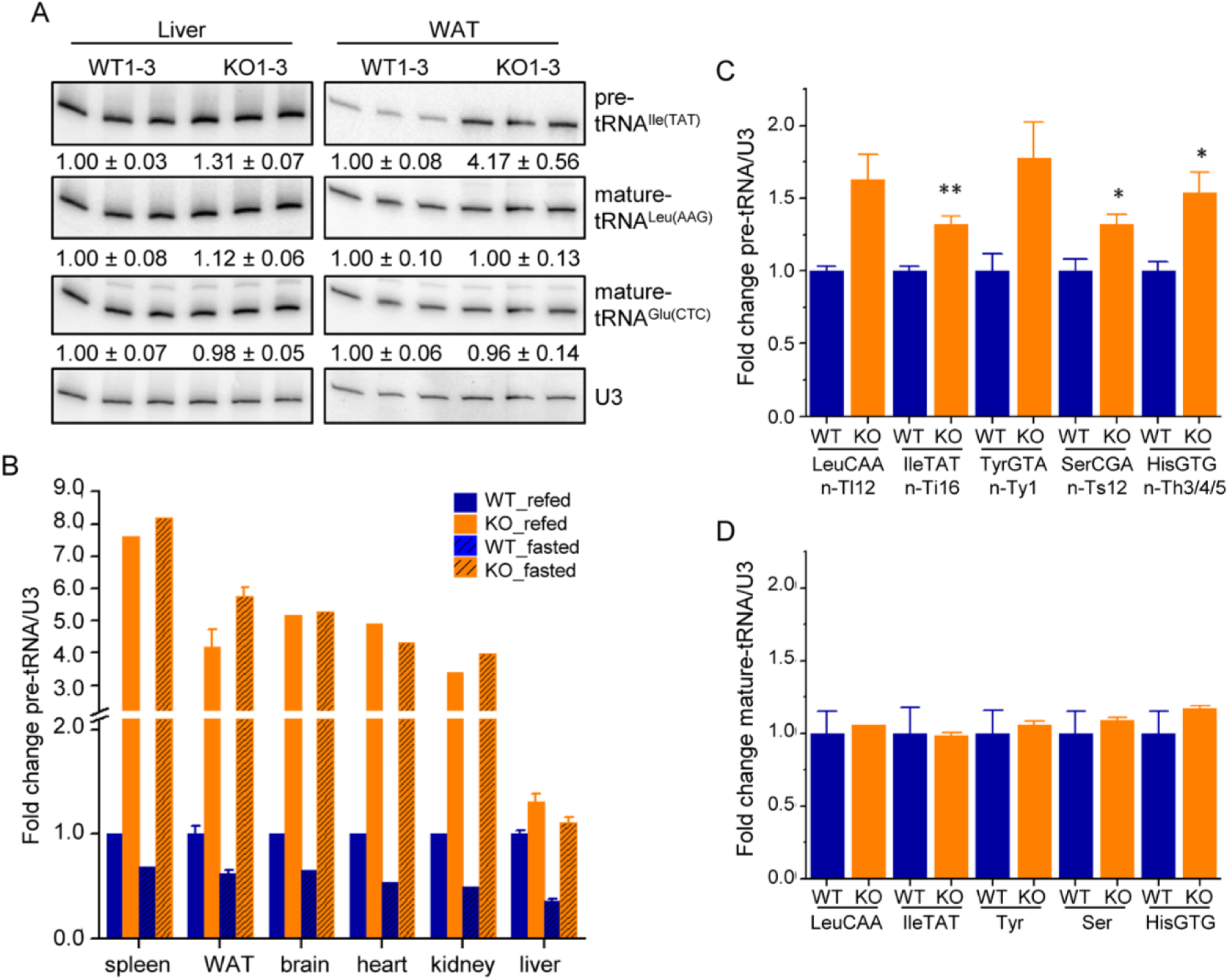
*Maf1*^-/-^ mice have higher levels of pre-tRNAs in refed and fasted conditions. (A) Northern blot analysis of liver and WAT mature-length and precursor tRNAs in 4 hour refed conditions. The intensity of each band was quantified and the mean values and standard errors (SEM) for the WT and *Maf1*^-/-^ independent biological replicates are indicated below each panel. Small nucleolar RNA U3 was used as a loading control. Note that all blots represent contiguous panels from the same gels. The images have been cropped at the top and the bottom to frame the relevant region for presentation. (B) Northern blot analyses of pre-tRNA^Ile^-TAT (locus n-Ti16, see Table S1) in the indicated tissues from 4 hour refed or 16 hour fasted WT and *Maf1*^-/-^ mice. The fold change in pre-tRNA normalized to U3 snRNA was expressed relative to the mean WT refed value for each tissue. WT and *Maf1*^-/-^ are represented by blue and orange columns, respectively, with solid colors reporting refed values and solid colors with cross-hatching reporting fasted values. Northern blot analyses of normalized pre-tRNAs (C) and mature tRNAs (D) in refed liver samples (WT n=3, KO n=4) were expressed relative to the mean value for WT (± SEM). For panel C, P values (Student’s standard t-test) are: pre-tRNA^Leu^ 0.07; pre-tRNA^Ile^ 0.007; pre-tRNA^Tyr^ 0.052; pre-tRNA^Ser^ 0.03 and pre-tRNA^His^ 0.025 and are indicated as * < 0.05 and ** < 0.01. Abbreviated locus information is annotated underneath each pre-tRNA, as listed in Table S1, column C.

### Analysis of the Pol II transcriptome in *Maf1*^*-/-*^ liver

Several studies have reported that perturbing Maf1 function by overexpression or knockdown affects the expression of certain genes transcribed by RNA polymerase II (reviewed in ^13^). In contrast, RNA-seq analysis of WAT from fasted wild-type and *Maf1*^-/-^ mice did not identify any significantly affected genes ^11^. To examine this question in refed liver, we generated libraries for high throughput sequencing from multiple independent WT (n = 7) and *Maf1*^-/-^ (n = 5) samples. After normalization for sequencing depth, read count data were scaled using TMM normalization and estimates of mean-variance (limma voom) and technical variation (RUVSeq) were computed for differential expression analysis with limma (see Materials and Methods). Moderated t-tests were then performed on the expression profiles and a cut-off on both the adjusted p-value (< 0.01) and the fold change (> log2 |0.5|) was used to select differentially expressed genes. This procedure identified 12056 expressed genes, 937 of which met the criteria for differential expression (Figure 4A and 4C, Table S6). The vast majority of these genes (89%) were downregulated in the *Maf1*^-/-^ samples. This asymmetric distribution was even more extreme at higher stringency cut-offs; 98% of 204 differentially expressed genes were downregulated at adjusted p-values < 0.001 (Table S6). To determine whether differential gene expression was independent of the average level of expression of each gene, i.e. if genes with high or low levels of expression were affected similarly, a reverse cumulative frequency was computed for the set of 937 genes. The expression of these genes was modestly lower in the liver of *Maf1*^-/-^ mice regardless of whether they were expressed at low or high levels (Figure 4B). The genes that showed the highest difference in mRNA accumulation between WT and *Maf1*^-/-^ mice were, as expected, *Maf1* (>30-fold decrease) and, more surprisingly, *Gm35339* (>30-fold increase, Figure 4C and Table S6). *Gm35339* is located immediately downstream of *Maf1* and is predicted to encode a WD40 repeat protein of unknown function. Overexpression of this gene could conceivably contribute to some of the phenotypes observed in *Maf1*^-/-^ mice described below. However, a role for *Gm35339* in obesity resistance is unlikely since two MAF1-null lines bearing different indels created by a targeted zinc finger nuclease do not have elevated levels of *Gm35339* transcripts yet resist weight gain on a high fat diet like *Maf1*^-/-^ mice ^11,23^.

**Figure 4.**
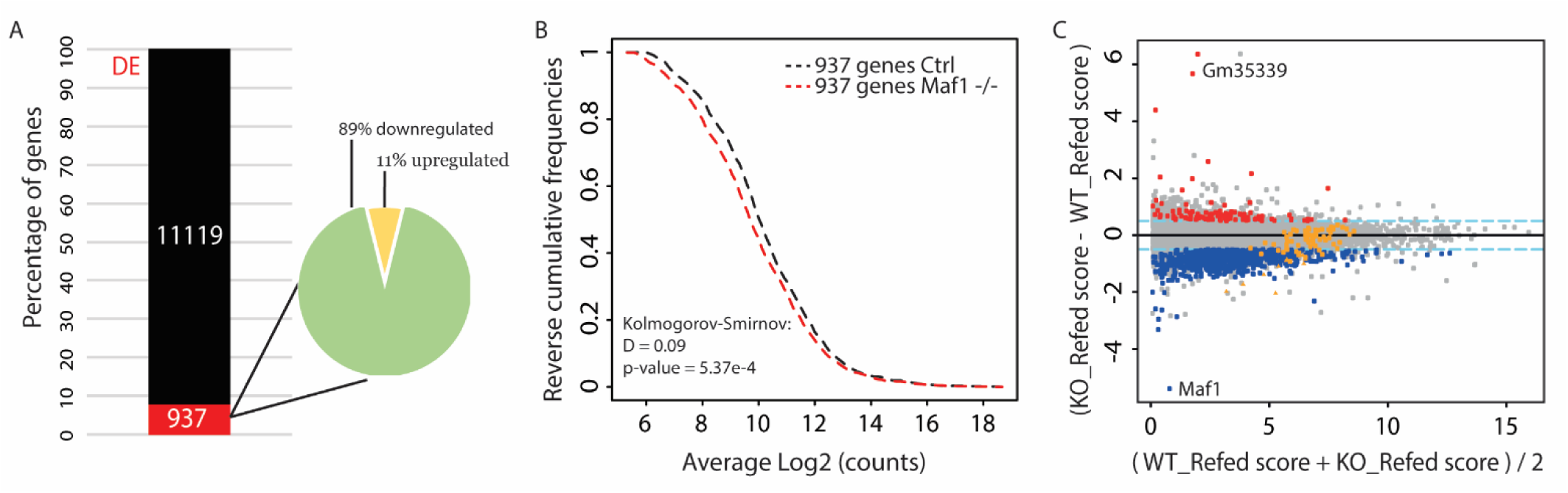
Changes to the Pol II transcriptome in fed liver from *Maf1*^-/-^ mice. (A) Representation of differentially expressed (DE, red) and unaffected (black) transcripts quantified by RNA-seq. The Venn diagram indicates the distribution of up- and down-regulated genes. (B) Relative reverse cumulative frequency plots are shown for the 937 differentially expressed genes in WT and *Maf1*^-/-^ samples. A Kolmogorov-Smirnov test was performed using the WT sample as the reference. (C) MvA plot representing the mean of the scores (log2(counts)) obtained in the WT and *Maf1*^-/-^ samples on the x-axis and the log2 fold change (*Maf1*^-/-^/WT) on the y-axis. Red and blue dots correspond to the set of 937 genes with adj. p-values < 0.01 and fold change > log2 |0.5| (dotted blue lines). Orange symbols show ribosomal protein genes that were significantly (triangles) or not significantly (dots) changed.

Gene ontology (GO) analysis of the differentially expressed transcripts identified a family of enriched GO bioprocess terms involving various aspects of lipid metabolism, storage and transport that relate to the lean phenotype of *Maf1*^-/-^ mice including previously reported changes in lipid droplets, liver triglyceride levels, lipolysis and fatty acid metabolism (Table S7) ^11,12^. Additionally, consistent with the bias towards gene down-regulation, numerous GO terms involving gene transcription and its negative regulation were recovered (Table S7). Another enriched GO term, translation, reflects the apparent down-regulation of genes encoding 10 subunits of the cytoplasmic ribosome (Figure 4C, orange triangles, Table S6). We also searched our gene list for protein coding genes (*Fasn, Acaca, TBP* and *PTEN*) whose expression has been reported to change upon *Maf1* overexpression and knockdown in *C. elegans* and/or in various cancer cell lines of hepatic origin ^15,17,24^, reviewed in ^13^. None of these genes were changed at our low stringency cutoff (Table S6). More recently, RNA-seq data has been reported following *Maf1* knockdown in mouse NIH3T3-L1 cells, a fibroblast-like cell line that can be chemically-induced to differentiate into adipocytes ^16^. Profiling of the undifferentiated NIH3T3-L1 cells identified 958 differentially expressed genes (575 up- and 383 down-regulated) at similar cutoffs (adj. P < 0.01 and a fold change >1.7) as our low stringency dataset. We note however that the absence of replicates in this study reduces the precision of the measurements and requires statistical tests to assume equal variance between the groups. A comparison with our set of 937 genes identified only 14 shared genes with concordant gene expression changes, significantly less than the 74 genes expected by random chance. We compiled the data from both studies to obtain 10852 shared genes and determined the threshold-free rank-rank hypergeometric overlap. This procedure is especially sensitive in identifying overlaps between weakly similar gene expression profiles ^25^. Despite this, no significant overlap was found between the top or bottom 2000 genes ranked by fold change or by sign-adjusted p values (Figure S2). Thus, the pattern of gene expression following knockout or knockdown of MAF1, along with MAF1 phenotypes, is surprisingly distinct in different cellular and nutritional contexts ^13^.

### Lack of MAF1 decreases translation in the liver of refed mice

In *C. elegans* and *D. melanogaster*, down-regulation of *Maf1* expression results in increased animal size ^14,15^ and increased translation ^14^. However, the reduced levels of several ribosomal protein mRNAs in our RNA-seq data suggested that translation might instead be negatively affected in the liver of *Maf1*^*-/-*^ mice. To assess global levels of translation, we first obtained polysome profiles by sucrose gradient sedimentation. Polysomal fractions from WT and *Maf1*^*-/-*^ mice were very similar under fasted conditions (Figure S3A). However, under refed conditions, we observed a reproducible reduction in polysomes in the *Maf1*^*-/-*^ samples, suggesting a reduced rate of protein synthesis (Figure 5A).

**Figure 5.**
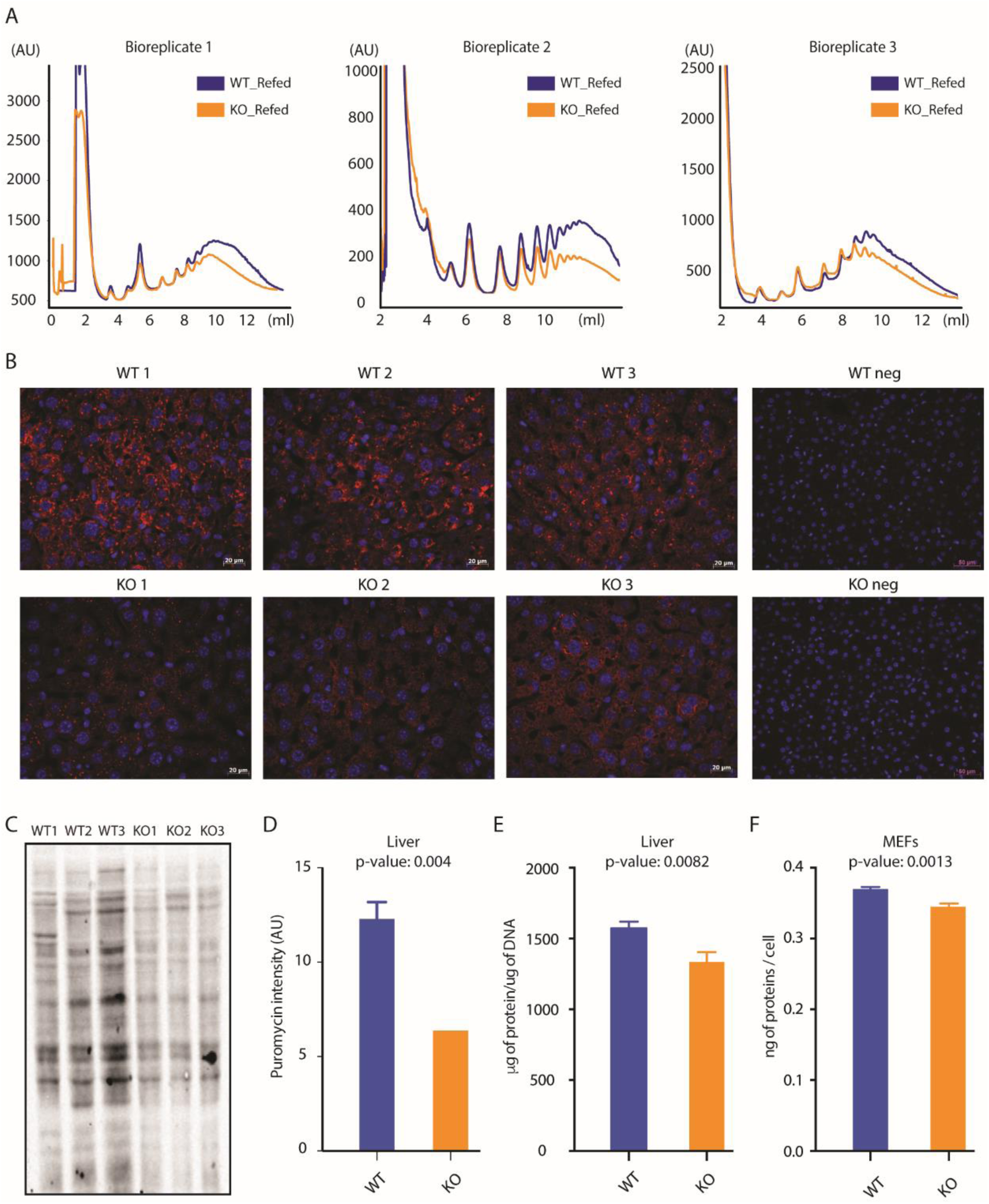
Reduced translation in the liver of fed *Maf1*^-/-^ mice. (A) Polysome profiles of liver samples from three WT (blue curves) and three *Maf1*^-/-^ (orange curves) refed mice. (B) Anti-puromycin immunocytochemistry of mouse liver. Mice were injected intraperitoneally with puromycin, livers were harvested after 30 min. and embedded in paraffin. Paraffin sections were incubated with an anti-puromycin antibody for detection of newly synthesized proteins (red signal). Nuclei were stained with DAPI (blue signal). The upper and lower panels on the right show negative controls from sham-injected mice. (C) Immunoblot of newly synthesized proteins in liver extracts from puromycin-injected mice. Samples were loaded according to DNA content and the blot was probed with an anti-puromycin antibody. (D) Quantification of the immunoblot in panel C. The signal obtained in each lane was normalized to total γ-tubulin, determined by re-probing the blot with an anti-γ-tubulin antibody. (Note that the difference was larger when the data were normalized for DNA content versus γ-tubulin). (E) Liver protein content normalized to DNA content. Protein and DNA were extracted and quantified from the same lysate for each of three WT and three *Maf1*^-/-^ livers. The ratios were plotted. (F) Protein content of mouse embryonic fibroblasts normalized to cell number. Proteins were extracted and quantified from three WT and three *Maf1*^-/-^ samples, each containing one million cells. Panels D-E report means and standard deviations.

Previous work by others suggests that translation efficiency of some classes of mRNAs vary during the diurnal cycle, with translation of mRNAs coding for components of the translation apparatus being highest during the night, two to six hours after the lights are turned off ^26^. To further characterize translation in *Maf1*^*-/-*^ mice, we pulse-labeled *ad libitum* fed mice two hours after dark for 30 min with low doses of puromycin, a structural analog of tyrosyl-tRNA, and monitored puromycin incorporation into nascent polypeptide chains ^27^. Puromycin immunohistochemistry showed a decreased cytoplasmic signal in *Maf1*^-/-^ liver (Figure 5B). Similarly, puromycin antibody probing of western blots showed that *Maf1*^-/-^ mouse liver incorporated as little as half the puromycin as WT tissue, whether the results were normalized for DNA content (Figure 5C) or total γ-tubulin signal (Figure 5D). Finally, total protein content normalized to DNA content was lower in *Maf1*^-/-^ liver compared to WT liver (Figure 5E), and total protein content normalized to cell number was lower in MEFs isolated from *Maf1*^-/-^ mice relative to MEFs from WT mice (Figure 5F). Thus, unlike what has been reported following *Maf1* knockdown in *Drosophila* ^14^, the complete absence of MAF1 results in decreased translation in the liver of fed mice.

### MAF1 function impacts translation efficiency in the liver of refed mice

To better characterize the change in translation, we performed ribosome profiling on liver from WT and *Maf1*^-/-^ mice fed *ad libitum* and sacrificed 2.5 hours after dark. Liver extracts from three mice in each group were spiked with a fixed amount of *Drosophila* S2 Schneider cell material as an internal control for sample-to-sample normalization, and to allow detection of global changes in translation. A comparison of the distribution of translation efficiency scores revealed a *Maf1*^-/-^ score curve shifted to the left relative to the WT score curve, indicating a significant general decrease in ribosome occupancy in the *Maf1*^-/-^ samples (Figure 6A, Table S8, Kolmogorov-Smirnov test = 2.2 × 10^−16^). An overall reduction (∼13% on average) in ribosome occupancy was also apparent from the fold change in translation efficiency scores compared on a gene-by-gene basis (Figure 6B, Table S8). The majority of mRNAs had lower translation efficiency (TE) scores in *Maf1*^*-/-*^ versus WT liver with 262 mRNAs being significantly lower in the knockout (Figure 6B, Table S8). A GO analysis of the genes in this group identified cytoplasmic ribosomal proteins as the most significant KEGG pathway and molecular function term (Figure S3B). Ribosomal proteins as well as other factors linked to protein synthesis are encoded by mRNAs which harbor a terminal oligopyrimidine (TOP) tract close to their 5’ end that serves to co-ordinate their translation ^28,29^. Interestingly, gene set enrichment analyses revealed high enrichment scores among the 2000 genes with the highest ribosome occupancy fold change for i) mRNAs coding for factors involved in translation, ii) TOP mRNAs, and iii) mRNAs coding for ribosomal proteins (Figure 6C).

**Figure 6.**
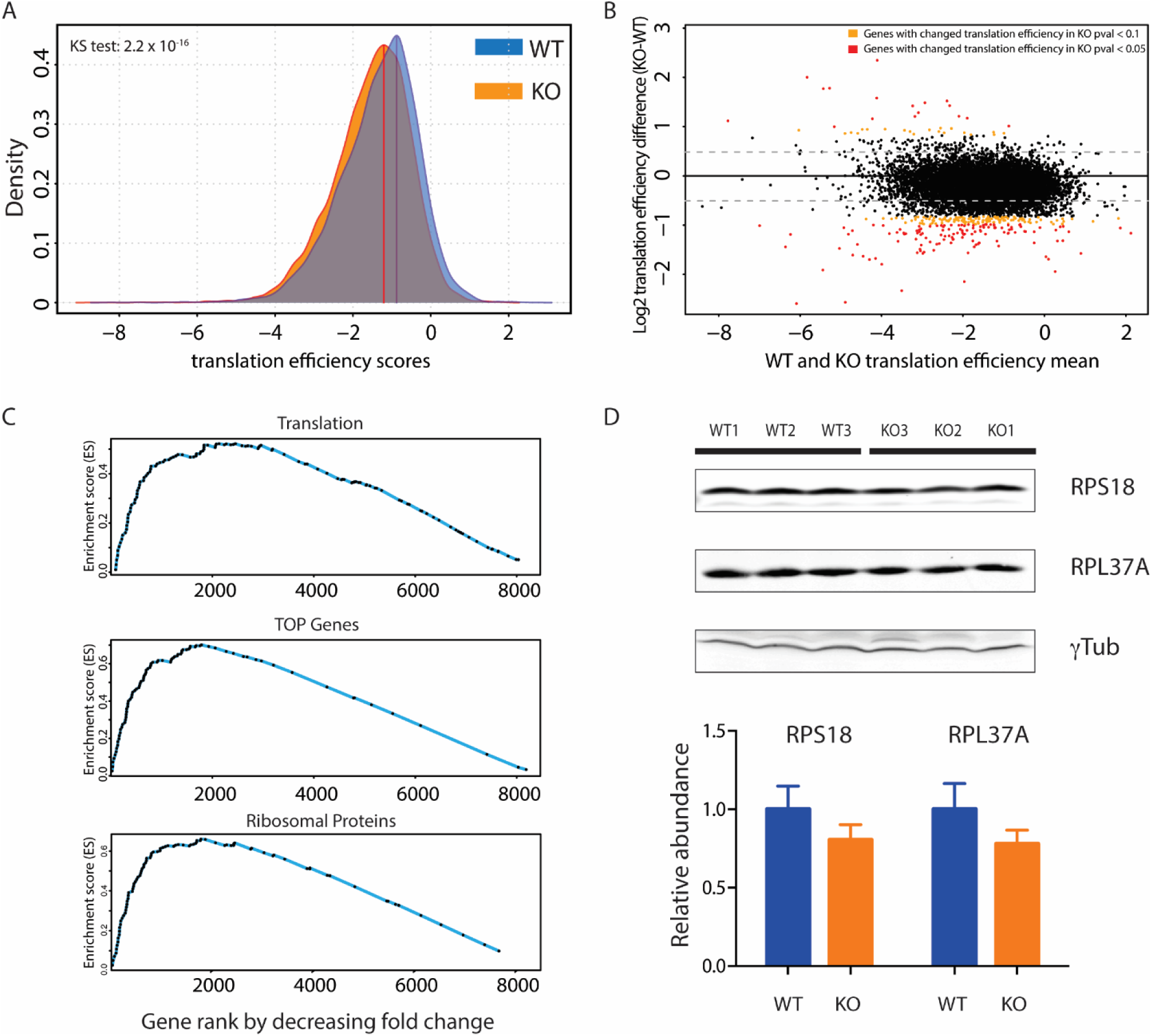
Ribosome profiling reveals global down-regulation of translation. (A) Kernel Density Estimation plot showing decreased ribosome occupancy on mRNAs in *Maf1*^-/-^livers. The x-axis shows translation efficiency scores (calculated as the average translation efficiency scores for all transcripts (splice site variants, alternative TSSs, etc.) derived from one gene, see Materials and Methods), the y-axis the score density. KS, Kolmogorov-Smirnov test. (B) MvA plot showing the average of translation efficiency scores in the WT and *Maf1*^-/-^ liver samples (x-axis) and the log2 fold change in *Maf1*^-/-^ versus WT samples (y-axis). Each dot represents the average score for all transcripts derived from one gene. The red and orange dots indicate score changes with adjusted p-values ≤ 0.05 and ≤0.1, respectively, as determined by limma. (C) GSEA analysis^56^ with the indicated gene set (genes encoding proteins involved in translation, TOP genes, and genes encoding ribosomal proteins). The x-axis shows genes ranked by decreasing fold change from left to right, the y-axis shows the running enrichment score for the gene set as the analysis progresses through the ranked list. (D) Immunoblots of liver lysates from WT or *Maf1*^-/-^ mice performed with antibodies directed against the indicated proteins. The lower panel shows the quantification of the signals normalized to γ-tubulin. Image acquisition was performed with the Odyssey Imaging System (LI-COR Biosciences) software (www.licor.com/bio/image-studio/), (version 2.1.10). The three panels are from the same gel, but image acquistion of RPS18 and RPL37A was performed on different channels.

Depending on the methods and statistical models used for analysis, translation efficiency scores determined from ribosome profiling experiments can be susceptible to high false discovery rates and low sensitivity ^30,31^. Thus, in addition using a generalized linear model for parameter estimation (limma, as above), we also analyzed the data using the negative binomial model of DESeq2, as implemented in Xtail ^31^. This program was shown to achieve high sensitivity while minimizing false positives in the analysis of simulated and published datasets including a study in which the translation of ribosomal proteins was perturbed by inhibition of mTOR signaling ^31^. At adjusted p value cutoffs of 0.05 and 0.1, Xtail was six- and nine-fold less sensitive, respectively, than limma in identifying mRNAs with differential translation efficiencies in *Maf1*^-/-^ versus WT samples (Tables S8 and S9). Nonetheless, four ribosomal protein mRNAs (Rps18, Rps20, Rpl11 and Rpl14) were listed among the 16 mRNAs whose translation efficiencies were reduced significantly in *Maf1*^-/-^ liver (adj. p values < 0.05, Table S9) and ribosome and translation were again the top scoring KEGG pathway and GO terms. The ability of MAF1 to affect the expression of ribosomal proteins was further supported by immunoblot analysis which revealed slightly (∼20%) reduced levels of large and small ribosomal subunit proteins, RPL37A and RPS18, in liver extracts from fed *Maf1*^-/-^ mice (Figure 6D).

## Discussion

Studies on the regulation of pol III transcription have typically examined acute responses of either yeast or mammalian cultured cells (e.g. to drugs, toxicants and nutrient withdrawal) that produce robust changes in activity. Relatively few studies have examined changes in pol III transcription that reflect physiological responses, such as feeding and fasting, in a mammalian organism. Looking into this issue, we observe that as WT mice fast, there is a decrease in pol III occupancy at loci that have low to intermediate occupancies in the refed condition, but essentially no change at loci with the highest occupancies, which tend to remain highly occupied after fasting (Figure 1). Indeed, pol III occupancy of most loci (265 out of 383 loci occupied in either the fasted or refed state) is not significantly affected during an 8 hour fast. Thus, fasting for 8 hours, a period in which metabolism switches from using mostly glucose as an energy source to using mostly lipids, does not constitute a substantial stress with respect to pol III transcription. This physiological control of pol III transcription in mice is distinctly different from the response of cultured human cells to serum starvation where the vast majority of pol III loci are subject to rapid MAF1-dependent repression ^9^.

A very different physiological response occurs in liver after a two-thirds partial hepatectomy as hepatocytes synchronously initiate mitotic growth to regenerate the organ. The recent analysis of pol III occupancy under these conditions allowed us to compare proliferative and nutritional responses. Our findings further support the concept of stable and changing populations of active pol III genes ^9,18^. Common sets of genes in both groups were readily identified. The changing population was notable because pol III occupancy of these genes can be driven up or down depending on the physiological signal. However, the observed changes in pol III occupancy in response to proliferation and nutrient signaling are more complex than a two state (changing versus stable) model since distinct sets of genes were identified that are preferentially responsive to one type of signal versus the other. These observations raise important questions for future studies. Why are the stable genes resistant to change? As described below, these genes are clearly controlled by MAF1, a direct mTORC1 substrate in mammals ^10^, so how do they resist change in response to fasting when mTORC1 activity is diminished? What mechanisms are responsible for the signal-dependent differential in pol III occupancy? Answers to these questions are likely to be functionally important given that the composition of the tRNA population is dynamic and can impact gene expression at multiple levels ^10,32,33^.

Our results also revealed that in the absence of MAF1, pol III occupancy in the liver and the levels of precursor tRNA in the liver and all other organs tested were higher than in WT mice, both in refed and fasted conditions. This changes our understanding of MAF1 from that of a repressor active only in response to stress or nutrient limitation to that of a repressor that keeps pol III output in check in different metabolic states, notably when energy derives mostly from glucose or lipids. Of note, in cultured cells, RNAi-mediated depletion of MAF1 results in higher pol III occupancy and transcription not only under serum-starved conditions, but also under serum-replete conditions ^9^. Thus, in both proliferating mammalian cells and in a differentiated mouse tissue, MAF1 functions as a chronic repressor whose activity can be modulated depending on the conditions.

We observed that pol III occupancy in the liver of *Maf1*^-/-^ mice is dramatically lower after an eight hour fast than after four hours of refeeding. Thus, MAF1 is not required for pol III to dissociate from the DNA in response to fasting. Since it is known that MAF1 bound to elongating pol III does not inhibit nucleotide polymerization or facilitated recycling ^34,35^, the results are consistent with a model of repression in which pol III must dissociate from the DNA after termination in order to be inhibited by MAF1 ^10^. Studies in yeast suggest that the switch between facilitated recycling and dissociation is modulated by nutrient and stress signaling to pol III and the initiation factor TFIIIB ^36,37^. In this system, pro-growth signaling leads to phosphorylation of Bdp1 and Rpc82 whereas under repressing conditions these modifications are lost and phosphorylation is acquired on Rpc53. The phosphoregulation of these proteins together is thought to promote pol III recruitment and facilitated recycling in favorable conditions and to promote pol III dissociation to enable repression by Maf1 when limiting nutrients or stress is encountered. Thus, our findings in mouse liver are consistent with the idea that like in yeast, MAF1-independent mechanisms promote pol III recruitment and dissociation in response to changing conditions.

Additional mechanistic insight is suggested by the fact that for the majority of pol III loci, the increase in pol III occupancy in *Maf1*^*-/-*^ versus WT liver is essentially the same for genes of low, intermediate or high occupancy. Linear regressions of significantly changing genes (red dots in Figures 2B and 2C) produce lines with slopes close to zero. This behavior is notably different from the physiological responses of stable and changing genes in WT tissue (Figure 1). Thus, MAF1 contributes to the differential response of these genes.

The greater magnitude of effect of *Maf1*^*-/-*^ on pol III occupancy over WT in refed versus fasted liver (Figure 2B and 2C) correlates with other changes under these conditions. Whole body energy expenditure is higher and metabolic changes in the liver involving amino acids and fatty acids are more prominent in the fed state versus the fasted state ^11,12^. Thus, the results are consistent with the idea that derepression of pol III transcription during all hours of the day, regardless of the feeding-fasting cycle, contributes to energy expenditure, with a larger share of that expenditure coming in the fed state. The wasteful use of this energy, which we previously proposed is driven by a futile RNA cycle ^11,12^, also seems to be enhanced in refed versus fasted liver: Higher levels of pol III occupancy in this condition are not accompanied by correspondingly large increases in precursor tRNAs in the liver (although large increases were observed in other tissues). This suggests that, reminiscent of *S. cerevisiae* where as much as 50% of pre-tRNA is degraded by the exosome before maturation ^38^, there is a high turnover of precursor tRNAs in fed *Maf1*^*-/-*^ liver. Importantly, mature tRNA levels in refed liver, as in the fasted state ^11^, remain largely unchanged through yet to be defined tRNA homeostatic mechanisms.

Pol II transcriptome profiling of liver from refed *Maf1*^*-/-*^ and WT mice identified a set of ∼1000 differentially expressed genes. A similar number of differentially expressed genes were reported in undifferentiated NIH3T3-L1 cells following MAF1 knockdown ^16^. However, the two profiles contained few overlapping genes and were not similar by a rank-rank hypergeometric test. In the liver of refed *Maf1*^*-/-*^ mice we identified ribosomal protein mRNAs as an enriched group of down-regulated genes. Interestingly, ribosomal protein genes were also enriched among mRNAs with reduced translational efficiency. These results correlate with a small overall reduction in global translation that is specific to the refed state, based on polysome profiles (Figures 4A and S3A), and with reduced incorporation of puromycin into nascent liver polypeptides. Thus, reduced levels of some ribosomal proteins resulting from lower mRNA levels and/or inefficient translation may reduce the total number of ribosomes for translation. This effect in mice was surprising since the opposite result was seen in *D. melanogaster.* Clearly, different mechanisms must be operating in mouse liver than those noted in *Drosophila* ^14^. Based on a variety of studies, MAF1 loss could affect transcription of some ribosomal protein genes ^39^ or reduce specific ribosomal protein mRNA levels through posttranscriptional mechanisms, e.g. by Argonaute-containing silencing complexes targeted by tRNA-derived fragments (tRFs) ^40,41^. We previously examined possible effects of the MAF1 loss on the synthesis of the large rRNAs and found no effect ^11^. Further, we did not observe downregulation of mTOR activity in liver from *Maf1*^*-/-*^ mice collected under various conditions as determined by relative phosphorylation levels of mTOR targets S6K, RPS6, and 4EBP1 (data not shown). Potential mechanisms affecting translation are many and may include tRF-mediated changes in mRNA structure ^42^, as well as impairment of mRNA decoding or triggering of an unfolded protein response ^43,44^ as a result of tRNA hypomodification ^45^. Our RNA-seq analysis of *Maf1*^-/-^ liver did not, however, indicate activation of the UPR. At any rate, the observed decreased in overall translation, which is consistent with the smaller size of *Maf1*^-/-^ mice ^11^, emphasizes how less MAF1 protein, which might be expected to lead to more tRNAs and thus more translation, as indeed it does in the fly, can have complex effects in the context of a mouse completely lacking MAF1.

## Materials and Methods

### Animals

All experiments involving mice were performed using protocols approved by the Veterinary Office of the Canton of Vaud (SCA-EXPANIM, Service de la Consommation et des Affaires Vétérinaires, Epalinges, Switzerland) in accordance with the Federal Swiss Veterinary Office guidelines or the Institutional Animal Care and Use Committee (IACUC) of the Albert Einstein College of Medicine. Mice were kept in a 12 hour day-12 hour night cycle, with light turned on at 7AM and turned off at 7PM. They were fed *ad libitum* on a chow diet except as specified below.

### ChIP-seq experiments

Mice at 12-14 weeks of age were fasted for 8 hours from 4PM to midnight and sacrificed, or fasted for 8 hours and refed for 4 hours, from midnight to 4AM, and sacrificed. Perfused livers were homogenized in 4 mL of PBS containing 1% of formaldehyde and left in the same buffer for cross-linking for 10 minutes. Nuclei were isolated as described in ^46^. Nuclear lysis was performed in 1.2 mL of 50 mM Tris-HCl (pH 8.1), 10 mM EDTA, 1% SDS, 50 μg/mL PMSF, 1 μg/mL Leupeptin. The nuclear lysate was then supplemented with 0.92 mL of 20 mM Tris-HCl (pH 8.1), 150 mM NaCl, 2 mM EDTA, 1% Triton X-100, 0.01% SDS, 50 μg/mL PMSF, 1 μg/mL Leupeptin and sonicated with a Branson SLPe sonicator for 10 cycles of 10 s at 50% amplitude, resulting in an average fragment size of 300 to 1000 bp. After each sonication cycle, the chromatin was kept in an ice-cold bath during 20s. Chromatin samples from three mice were pooled, de-cross-linked, and an aliquot was extracted for DNA quantification. Human HeLa cell chromatin was prepared as described in ^21^.

ChIPs were performed with 30.8 μg of total chromatin DNA containing 2.5% of human spike-in chromatin ^22^, and 10 μL of serum from a rabbit immunized with a peptide 100% conserved between human and mouse POLR3D (CS681 antibody, C-terminal peptide CSPDFESLLDHKHR, custom made for us by Covance) ^47^. This antibody has been used extensively for ChIP-seq experiments, both in human and mouse cells ^21,48,49^. The ChIPs were performed as follows. The chromatin samples were incubated with the antibodies overnight at 4°C. The next day, 20 μL of protein A sepharose beads (CL4B GE Healthcare) was added and the samples were further incubated for 3 h. The beads were next washed once with 20 mM Tris-HCl (pH 8.1), 50 mM NaCl, 2 mM EDTA, 1% Triton X-100, 0.1% SDS, twice with 10 mM Tris-HCl (pH 8.1), 250 mM LiCl, 1 mM EDTA, 1% NP-40, 1% deoxycholic acid, and twice with TE buffer 1X (10 mM Tris-HCl (pH 7.5), 1 mM EDTA). Subsequently, bound material was eluted from the beads in 300 μL of elution buffer (100 mM NaHCO_3_, 1% SDS), treated with RNase A (final concentration 8 μg/mL) over 6 h at 65°C and then with proteinase K (final concentration 345 μg/mL) overnight at 45°C. The next day, the samples were purified with PCR clean-up kit from MACHEREY-NAGEL and eluted in 50 μL of elution buffer.

DNA (10 ng) of from each ChIP was used to prepare sequencing libraries according to the Illumina ChIP-Seq DNA Sample Prep Kit protocol (Illumina; San Diego, California, USA; cat. no IP-102-1001), except that size selection of the samples was performed after, rather than before, library amplification. Sequencing libraries were loaded onto one lane of a HiSeq 2000 flow cell and sequenced at 100 cycles. For each condition we sequenced input chromatin and the corresponding ChIP sample(s). Data analysis was performed as described previously using spiked-in human chromatin for sample-to-sample normalization ^22^. Occupancy scores we calculated for 672 annotated pol III genes and other pol III occupied loci and genes were defined as “not occupied” or “occupied” based on a cut-off calculated as before ^49^.

### Northern Blot analysis

Mice fed *ad libitum* on a breeder chow diet were fasted overnight or fasted overnight and refed for 4 hours before sacrifice. Tissues were flash frozen in liquid N_2_ and stored at -80°C. Samples (50-100 mg) were homogenized in TRIzol (Thermo Fisher Scientific) and RNA was purified according to the manufacturer’s instructions. RNA was precipitated twice, quantified, and resolved by denaturing polyacrylamide electrophoresis before electrophoretic transfer to nylon membranes (GE Healthcare Amersham Hybond-N^+^) and hybridization with [^32^P]-end labelled oligonucleotide probes at 42°C. Hybridization signals detected by phosphorimaging were quantified and normalized to U3 snRNA to compare expression in WT and *Maf1*^-/-^ samples. The probe sequences were:

mature tRNA^Glu^-CTC 5’-CGCCGAATCCTAACCACTAG-3’;

mature-tRNA^Leu^-CAA 5’-CTCCATTCGGAGACCAGAA-3’;

pre-tRNALeu-CAA 5’-CCCGTAGGTAAGGCCTTGTCA-3’;

mature tRNA^Ile^-TAT 5’-TCCAGGTGAGGATCGAACTC-3’;

pre-tRNA^Ile^-TAT 5’-ATCGCTTACGCCTAGCACTG-3’;

mature tRNA^Tyr^-GTA 5’-GAGTCGAACCAGCGACCTA-3’;

pre-tRNA^Tyr^-GTA 5’-AAGGATGTCTTCTAACGGGGA-3’;

mature tRNA^His^-GTG 5’-CTATACGATCACGGCGAGCTACC-3’;

pre-tRNA^His^-GTG 5’-CCCTTGCCGTACCATTGTTGCCAC-3’;

mature tRNA^Ser^-CGA 5’-ACTCGGCCATCACAGCTAACC-3’;

pre-tRNA^Ser^-CGA 5’-GCTGTGTCTGAAGCTTTCTTACGCTGT-3’.

U3 snRNA:5’-GGAGGGAAGAACGATCATCA-3’.

Probes for pre-tRNAs recognize the unique introns of specific genes for tRNA^Leu^-CCA, tRNA^Ile^-TAT and tRNA^Tyr^-GTA and a unique 3-’trailer sequence for tRNA^Ser^-CGA. The 3’-trailer probe for tRNA^His^-GTG recognizes the transcripts from 3 genes.

### RNA-seq

Mice were fed a chow diet *ad libitum* until 12-14 weeks of age. Prior to the sacrifice, mice were fasted overnight and then refed for four hours. Livers perfused with phosphate buffered saline (10-30 mg pieces) were added to 1 ml of TRIzol (Life Technologies), mechanically homogenized, and RNA was extracted according to the manufacturer’s instructions. Samples were processed from seven wild-type mice and five *Maf1*^-/-^ mice. RNA concentration was determined with a NanoDrop ND-1000 spectrophotometer and RNA quality was assessed with the Fragment Analyzer (Agilent Technologies). Libraries were prepared with the TruSeq Stranded Total RNA Ribo-Zero Globin kit (Illumina) according to the manufacturer’s instructions. Sequencing libraries were loaded onto one lane of a HiSeq 2000 flow cell and sequenced at 100 cycles.

For analysis of RNA-seq data, purity-filtered reads were trimmed (cutadapt version 1.8) and reads matching to ribosomal RNA sequences were removed (FastQ Screen version 0.9.3). The remaining reads were further filtered for low complexity (reaper version 15-065)^50^ and then aligned against the *Mus musculus* GRCm38.86 genome using STAR (version 2.5.2b)^51^. The quality of the RNA-seq data alignment was assessed using RSeQC (version 2.3.7)^52^ and the number of read counts per gene locus was summarized with htseq-count (version 0.6.1)^53^. Library sizes were scaled using TMM normalization (edgeR version 3.14.0)^54^, expressed genes were selected with voom, unwanted variation factors were then calculated on the expressed genes with RUVSeq in edgeR, normalization and unwanted variation factors were then used in a the limma differential expression analysis. Differential expression was computed with limma ^55^ using the Benjamini-Hochberg procedure to control the false discovery rate.

### Sucrose gradient centrifugation of polysomes

Liver samples (50-100 mg) from 22-24 week old mice fasted or fasted and refed as described above for the ChIP-seq experiments were dounce homogenized in 20 mM Tris-HCl pH 7.4, 150 mM NaCl, 5 mM MgCl_2_, 1 mM DTT, 100 μg/mL cycloheximide, 1% Triton with 25 units/mL TURBO DNase 1, 0.2 units/mL RNasin, 1 mg/mL heparin, and cOmplete Protease Inhibitor Cocktail. Homogenates containing 50 μg of total RNA were loaded on 15%-60% sucrose gradients and centrifuged at 35000 rpm, 4°C for 4 h in an SW 40 rotor. Absorbance profiles at 260 nm were determined during gradient fractionation using an ÄKTA Purifier 10 (GE Healthcare).

### Anti-puromycin Immunohistochemistry

Puromycin (40 nmol/g body weight) was injected intraperitoneally into twelve-week old mice fed *ad libitum* at 9 PM, i.e. two hours after the lights were turned off (ZT14). Livers were harvested after 30 min. Samples for Western blotting were prepared by dounce homogenization in RIPA buffer, then normalized for DNA content and resolved on a denaturing 10% polyacrylamide gel. Proteins were transferred onto a Protran nitrocellulose membrane and probed with antibody clone 12D10 against puromycin (Millipore MABE343). Subsequently, the blot was redetected with an antibody against γ-tubulin (Santa Cruz, antibody B-12).

For immunostaining experiments, paraffin-embedded liver sections were prepared from puromycin-injected mice after fixation of the tissue for 8 hours in PBS-buffered 4% paraformaldehyde. Rehydrated sections were incubated with 0.01 M citrate buffer, pH 6, for 10 min and then washed extensively in PBS. To prevent non-specific binding, proteins were blocked with 1% normal goat serum (30 min at room temperature) followed by a goat anti-mouse IgG2a antibody (Abcam, ab98694). After additional blocking with normal goat serum, the sections were incubated at 4°C overnight with anti-puromycin antibody and visualized using an Alex Fluor 568 conjugated goat anti-mouse IgG (Thermo Fisher, A11019). Nuclei were stained with DAPI (Sigma-Aldrich, D9542) at a dilution of 1/5000.

### Genomic DNA extraction

A small piece of liver was homogenized in 50 mM Tris-HCl pH 8.0, 100 mM EDTA, 0.125% SDS, 0.8 mg/ml proteinase K and then incubated overnight at 55°C. After sequential phenol/chloroform extractions, the DNA was precipitated and resuspended for fluorometric assessment of concentration using Qubit technology (Invitrogen).

### Ribosome profiling

Livers collected from 12 week-old mice fed *ad libitum*, two hours after the lights were turned off, were directly lysed in the polysome buffer described above and kept on ice for 10 min. Samples were centrifuged at 3000 rpm for 3 min and the supernatants were used for library preparation. Similar material extracted from Drosophila S2 cells was used as a spiking control and added to the mouse material at a ratio of 1/15. Thirty ODs of the spiked material was used for library preparation. Libraries for Ribo-seq and total RNA-seq were prepared in parallel according to the Illumina TruSeq Ribo Profile (Mammalian) protocol. The Truseq Ribo Profile Nuclease was replaced by the RNase I from Ambion (cat no AM2295) to generate the ribosome footprints.

Sequence tags from RNA-seq and from ribosome footprints performed in parallel from the same sample were aligned to both the *Drosophila melanogaster* BDGP Release 6 plus ISO1 MT/dm6 and *Mus musculus* NCBI37/mm9 genome assemblies, common tags were eliminated, the remaining tags were then aligned onto the dm6 and mm9 coding sequences (CDSs) (in a first step, without any mismatch allowed, and in a second step with up to one mismatch allowed), then all aligned tags were remapped and their position on genes recorded. Tags aligning to regions encoding rRNA or scRNAs were eliminated, and the remaining aligned tags were normalized i) to the total number of aligned sequence tags in each experiment, and ii) with TMM from edgeR ^54^. A correction factor calculated from the signal of the *Drosophila* S2 Schneider cell spike material was then applied to the normalized tag numbers (log2) to obtain scores for individual transcript types. The average of the scores for all transcripts derived from a single gene (with different TSSs, splicing patterns, or poly A addition sites) was then calculated to obtain the RNA-seq-normalized tag numbers (RNA-seq-NTC) and the ribosome footprint normalized tag counts (RF-NTC) listed in Table S8. Translation efficiency scores and p-values were determined from these values using the programs limma and Xtail ^31,55^. Antibodies to RPS18 (ab91293) and RPL37 (ab194669) were from Abcam.

## Data Availability

The ChIP-seq, RNA-seq and ribosome profiling data in this study have been submitted to the NCBI Gene Expression Omnibus (GEO; http://www.ncbi.nlm.nih.gov/geo) under the accession number GSE104535.

## *Ack*nowledgements

We thank Henrik Kaessman and Angelica Liechti for providing us with Drosophila S2 cell extracts, and Peggy Janich and David Gatfield for help with the ribosome profiling experiment. We also thank Pascal Cousin for help with experiments and analyses, and Lluis Fajas Coll and his group for valuable discussions. This work was funded by the University of Lausanne and Swiss National Foundation grants (31003A_169233 and CRSI33_125230) to NH and a National Institutes of Health grant (GM120358) to IW.

## Author Information

### Contributions

N.B designed and performed experiments, analysed data and wrote the paper. V.P performed the bioinformatics analysis. R.M, G.W, and F.M performed experiments. C.M performed the immunocytochemistry. I.W and N.H designed experiments, analysed data and wrote the paper.

## Ethics declarations

### Competing Interests

The authors declare no competing interests.

## Supplementary Information

Supplementary information.

## Notes

#### Summary of Updates

This version of the manuscript contains minor changes to the text and improved labeling of some figures.

