## Supplemental Information for "MAF1 is a Chronic Repressor of RNA Polymerase III Transcription in the Mouse"

**Chronic repression by MAF1 supports futile RNA cycling as a mechanism for obesity resistance**

**Supplementary Information**

**Figure S1. Analysis of Pol III occupancy scores.**

**Figure S2. Rank-rank hypergeometric overlap heatmaps.**

**Figure S3. Polysome profiles and bioinformatic analysis of ribosome profiling data.**

**Table S1. Pol III occupancy of the loci examined in this work. Related to Figures 1 and 2.**

**Table S2. Pol III Loci above the cut-off in at least one condition shown in Figure 1B.**

**Table S3. Pol III Loci above the cut-off in at least one condition shown in Figure 2B.**

**Table S4. Pol III Loci above the cut-off in at least one condition shown in Figure 2C.**

**Table S5. Pol III Loci above the cut-off in at least one condition shown in Figure 2D.**

**Table S6. RNA-seq analysis of fed liver from wild-type and *Maf1*$^{-/-}$ mice.**

**Table S7. GO bioprocess enrichment of differentially expressed genes in fed liver of wild-type *Maf1*$^{-/-}$ mice.**

**Table S8. Ribosome footprinting of liver samples from wild-type *Maf1*$^{-/-}$ mice.**

**Table S9. Xtail analysis of ribosome footprinting data.**

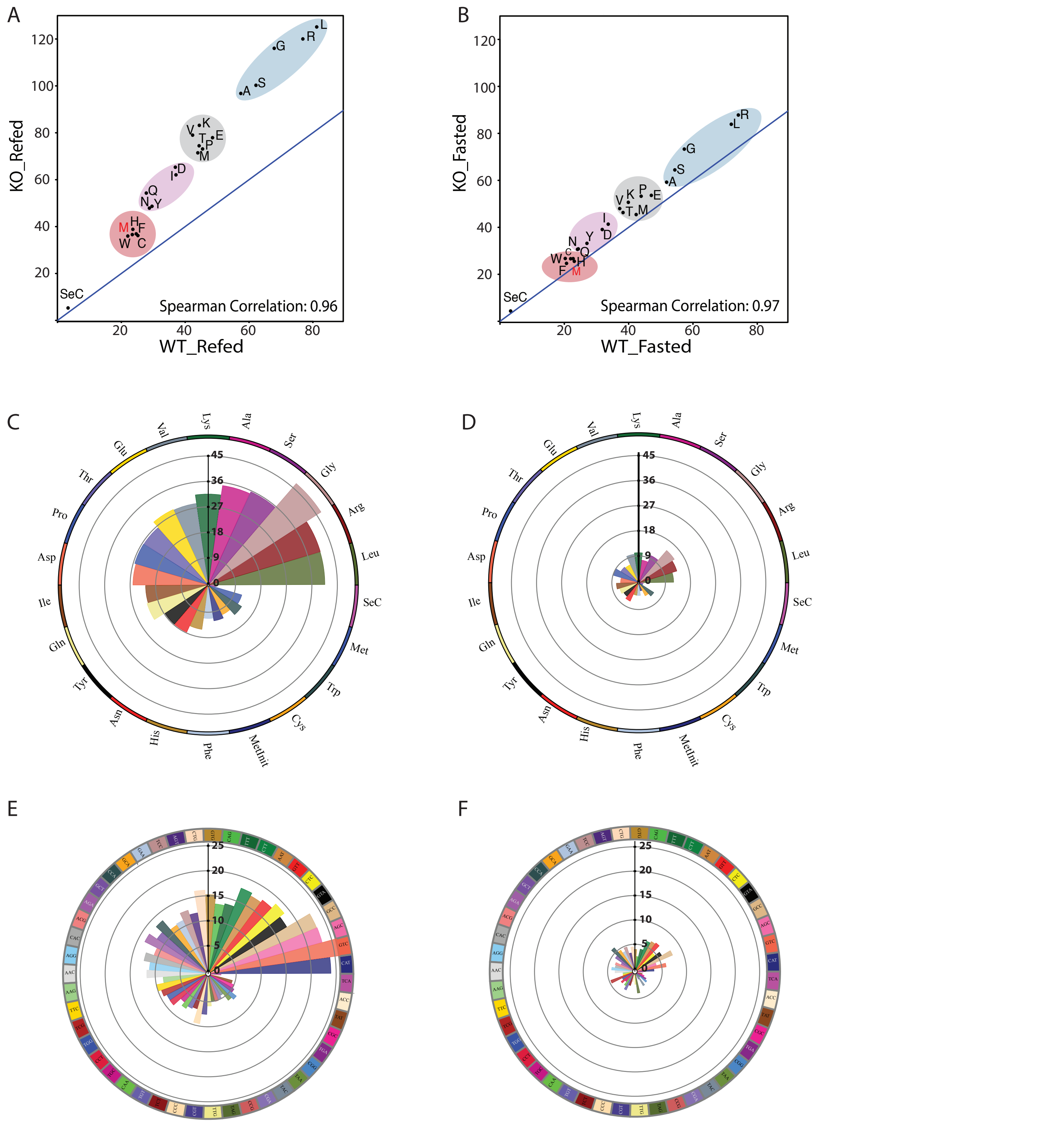


**Figure S1.** (A) Spearman rank correlations of pol III occupancy scores cumulated per isotype in WT and Maf1$^{-/-}$ mice in the refed condition. Twenty-one tRNA gene isotypes (tRNA$^{SeC}$ gene not included and tRNAi$^{Met}$ genes were considered separately from tRNA$^{Met}$ genes) were clustered in four different quantiles (blue, grey, purple and pink ovals) according to their pol III occupancy. (B) As in panel A but for mice in the fasted condition. (C) Circular plot indicating the ratio of cumulated pol III occupancy scores between Maf1$^{-/-}$ and WT samples for each tRNA isotype in the refed condition. Isotypes were ordered according to cumulated scores in the WT sample. (D) As in panel C but for the 8 hour fasted condition. (E) Circular plot indicating the ratio of cumulated pol III occupancy scores between Maf1$^{-/-}$ and WT samples for each tRNA isoacceptor in the refed condition. Isoacceptors were ordered according to cumulated scores in the WT sample. (F) As in panel E but for the 8 hour fasted condition.

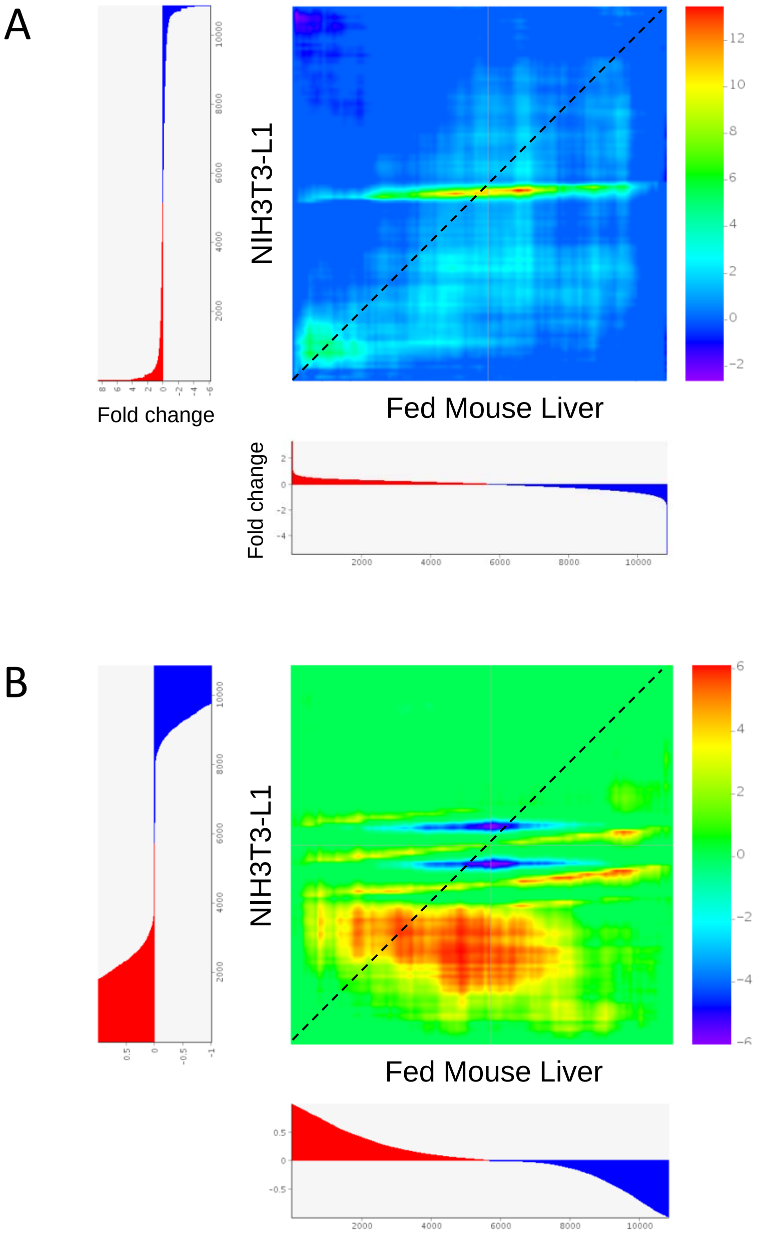


**Figure S2.** Rank-rank hypergeometric overlap heatmaps. The effects of *Maf1* knockout or knockdown on gene expression are compared in mouse liver and undifferentiated NIH3T3-L1 cells, respectively. Overlaps were computed from a list of 10852 genes that were scored in both samples (this work Table S6 versus GEO dataset GSE113324 day zero samples before addition of differentiation cocktail). (A) Genes are ranked by fold change in gene expression. (B) Genes are ranked by adjusted log10 P values that are sign-adjusted depending on whether gene expression increased or decreased. In both panels, correlated gene expression patterns lie in proximity to the diagonal (dotted line) with upregulated genes in the bottom left quadrant and downregulated genes in the top right quadrant. The scale bar on the right shows the log10-transformed hypergeometric P values (Benjamini-Yekutieli corrected) with positive values indicating over-enrichment and negative values indicating under-enrichment. The metric values for each heatmap are plotted adjacent to the x- and y-axes and are colored red or blue to indicate increased or decreased values, respectively. Note that the highest hypergeometric P-values in both maps (red) correspond to gene ranks that are uncorrelated or correspond to fold changes or adjusted P values that are below thresholds for significance.

A

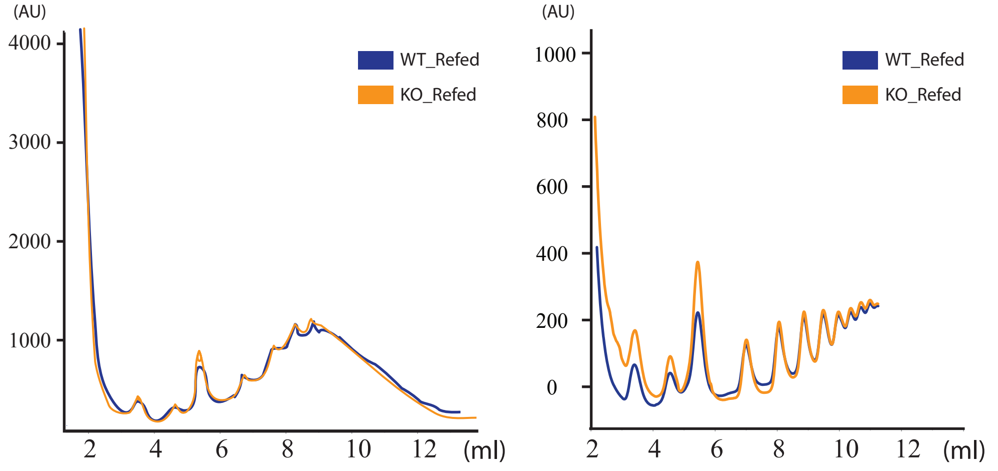


B

| KEGG pathways | Adj. Pvalue |
|---|---|
| Ribosome | $2.14 \times 10^{-05}$ |
| Ovarian steroidogenesis | 0.0032 |
| Proteoglycans in cancer | 0.0060 |
| Insulin secretion | 0.0138 |
| One carbon pool by folate | 0.0162 |
| Long term depression | 0.0167 |

| GO molecular functions | Adj. Pvalue |
|---|---|
| Structural constituent of ribosome | $7.16 \times 10^{-05}$ |
| Protein binding | 0.0029 |
| Transcription factor activity | 0.0058 |
| Profilin binding | 0.0059 |
| ATP-dependent microtubule motor activity | 0.0094 |
| Phosphoprotein phosphatase activity | 0.0122 |

**Figure S3.** A) Polysome profiles of liver samples from three WT (blue curves) and three Maf1$^{-/-}$ (orange curves) 22-24 week old fasted mice. (B) KEGG pathway (left) and GO molecular function (right) analysis for the genes showing lower translation efficiency (p-value <0.1) in the ribosome profiling experiment.
